# matchSCore: Matching Single-Cell Phenotypes Across Tools and Experiments

**DOI:** 10.1101/314831

**Authors:** Elisabetta Mereu, Giovanni Iacono, Amy Guillaumet-Adkins, Catia Moutinho, Giulia Lunazzi, Catarina P. Santos, Irene Miguel-Escalada, Jorge Ferrer, Francisco X. Real, Ivo Gut, Holger Heyn

## Abstract

Single-cell transcriptomics allows the identification of cellular types, subtypes and states through cell clustering. In this process, similar cells are grouped before determining co-expressed marker genes for phenotype inference. The performance of computational tools is directly associated to their marker identification accuracy, but the lack of an optimal solution challenges a systematic method comparison. Moreover, phenotypes from different studies are challenging to integrate, due to varying resolution, methodology and experimental design. In this work we introduce *matchSCore (https://github.com/elimereu/matchSCore)*, an approach to match cell populations fast across tools, experiments and technologies. We compared 14 computational methods and evaluated their accuracy in clustering and gene marker identification in simulated data sets. We further used *matchSCore* to project cell type identities across mouse and human cell atlas projects. Despite originating from different technologies, cell populations could be matched across data sets, allowing the assignment of clusters to reference maps and their annotation.

## Introduction

Single-cell RNA sequencing (scRNA-seq) is a powerful tool to quantify phenotype heterogeneity across individual cells. Recent discoveries of novel cell types and states provide the basis for more refined classifications of complex tissues and dynamic systems^1^. Rapid improvements of single-cell technologies and corresponding computational solutions allowed the processing of large sample sizes, resulting in the first cell atlases of tissues, organs and organisms^2–5^. Single-cell data analysis involves cell clustering to define biologically distinct types or to describe dynamic processes where cells transform in a continuum of states. Cells are grouped by transcriptional similarities before specific marker genes are identified. Both steps critically impact on the phenotyping resolution by providing population structures and their molecular characteristics. However, the analysis of single-cell transcriptomics data sets is challenged by the stochastic component of gene expression and technical sources of noise (e.g., dropout events and batch effects)^6, 7^. Further, complex subtype structure and large intra-group variability lead to highly variable gene expression distributions and fuzzy clusters. Hence, biological and technical factors can confound the cellular population structure, critically impacting on the performance of analytical tools and challenging the integration of data sets across studies and technologies.

Comparative work has been performed for differential expression analysis tools^8–10^, but approaches for clustering and gene marker identification have not been systematically tested. To date, cluster quality assessments were based on robustness and stability^11–13^, cohesion^11^ or the visual inspection of cluster separation^14^. Validation was performed through classification techniques, such as support vector machines^11, 15^ or supervised with established gene markers^16^. However, in the light of the attempts to generate comprehensive cellular atlases, cluster accuracy and marker identification become extremely important to explain the molecular basis underlying phenotype formation. It is the combined performance in cell clustering and gene marker identification that defines a method’s suitability to describe sample complexity. Moreover, the power to identify population gene markers strongly relates to the type and quality of the underlying data sets, challenging the integration of experiments from different studies and technologies. However, straightforward matching of future data sets into current efforts to produce reference cell atlases of organisms is strongly desired. In this regard, researchers should be enabled to compare their experimentally derived clusters with reference maps, to annotate cells and to determine perturbation effects, such as present in diseases. Recent computational tools enable data integration across experiments through normalization or the projection of cells into a reference maps^17–19^. However, these methods do not provide a straightforward solution for cell type annotation or use heuristic steps and distance quantifications that are suboptimal for the context of noisy single-cell gene expression data. To enable systematic comparison of computational tools and straightforward cross-study data integration, we introduce *matchSCore*, a Jaccard index based scoring system, to quantify clustering and marker accuracy in a combined score and to integrate cluster identities across different data sets. The approach allowed us to evaluate consistencies between simulated populations and predicted clusters from commonly used computational tools. Specifically, we performed simulation-based benchmarking of 14 single-cell phenotyping methods by measuring their agreement with a simulated optimal solution. The *matchSCores* could be tracked at different thresholds, thereby providing a trend of accuracy and indicating the sweet spots of the tested methods. Our comparative analysis will help users to make an informed decision to select tools tailored to their respective data sets and priorities. Next, we used the metric of *matchSCore* to combine results from single-cell RNA sequencing data sets from different studies. Specifically, we produced single-cell transcriptome profiles for bladder and pancreas tissues and projected our and publically available cell clusters onto a mouse or human organ reference atlas, respectively. The integration of tissue-matched data sets across different RNA sequencing techniques (full-length and 3’-digital counting methods) underlined the broad utility of our scoring system to annotate a given data set in the framework of a predefined reference.

## Results

We introduce *matchSCore* as metric to integrate cell type identities by combining clustering information with associated population markers. To derive a *matchSCore*, simulated groups of cells are matched to computational clusters to assign true-positive group labels. Then, predicted and simulated group markers are compared to determine consistencies (**Fig.1**). The matching of markers takes into account their group and cluster specificity (marker ranking), according to their fold-change (simulation) and p-value (tool), respectively. Highly specific genes (for a group or cluster) are top-ranked, while shared markers across groups (low-specific markers) are ranked at lower positions. Alternatively, clusters of a given single-cell RNA sequencing experiment are mapped to a reference data set to score cell identity similarities. In both contexts, the Jaccard index^20^ is the metric utilized to determine similarities across simulated or reference group markers and their predicted or experimentally derived counterparts, respectively. The Jaccard index represents the standard metric to assess the accuracy in object detections and image segmentations. In that context, accuracy is determined by the matching area over the total area of a ground truth and a tested image. Similarly, we used the Jaccard index to compare experimentally derived gene markers to an optimal solution or reference. The index allows features in multiple groups and thus provides an adequate measure for marker set similarities (including shared markers across groups). It is of note that the *matchSCore* tolerates imperfect clustering, but strictly penalizes low marker accuracy. With cell clustering being inevitably impacted by the incomplete and stochastic nature of single-cell data sets, marker genes are crucial to support computational clusters and to guide biological interpretation. We used *matchSCore* to benchmark clustering accuracy and the sensitivity to identify gene markers of computational tools with respect to an optimal (simulated) solution. Then we applied our scoring metric to match cell type identities of different organs across experiments and scRNA-seq methods.

**Figure 1.**
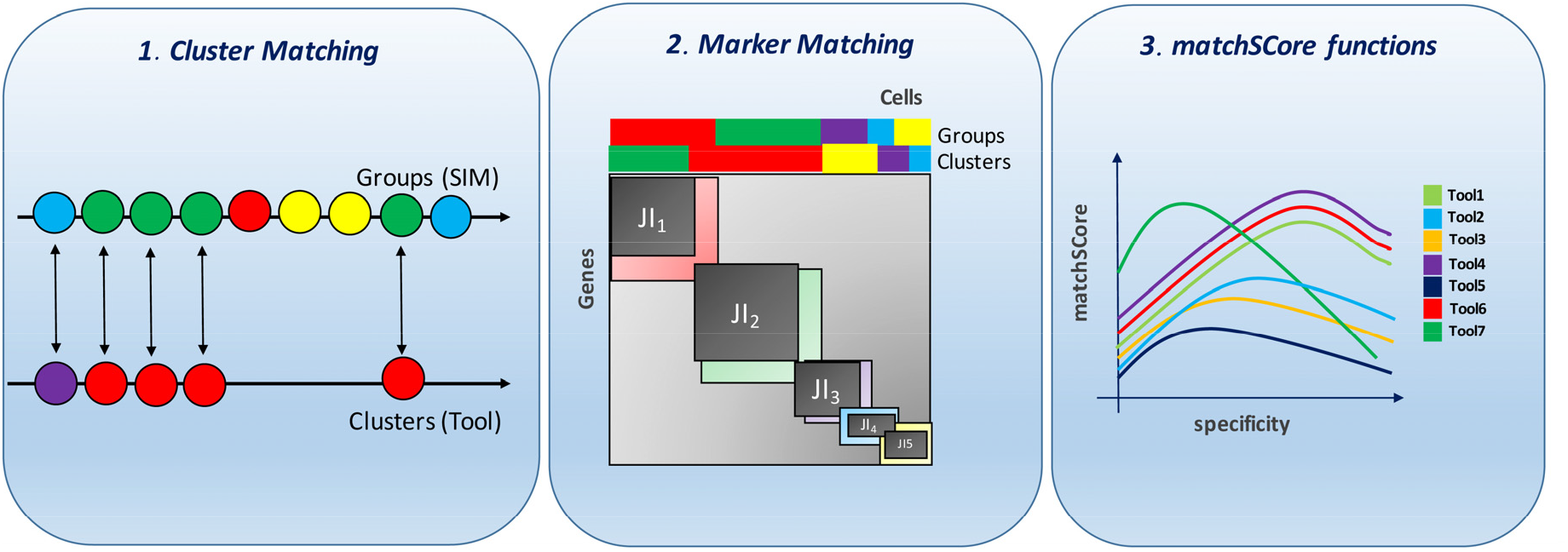
Schematic representation of the *matchSCore* metric to measure similarity and accuracy between computational clusters with relative cluster-specific genes and reference groups with true group markers. The metric includes a step to identify true group labels based on the highest match between simulated groups and clusters. Then the Jaccard index is computed across corresponding sets of markers. A *matchSCore* can be computed at different level of marker specificity to identify the performance peak of a tool.

### Benchmarking computational tools for cellular phenotyping

We simulated four levels of complexity for two common scenarios in single-cell experiments: a) a heterogeneous mixture of cell types and subtypes (S1) and b) a dynamic process with continuous cellular states (S2). The simulations were based on two reference data sets from which genes and cell numbers, gene expression mean, variance and library size were estimated (**Online Methods**). Specifically, we used scRNA-seq data from 2700 peripheral blood mononuclear cells, available from 10x Genomics (PBMC; S1), from which we simulated eight cell populations (including six with proportions of 0.1 and two of equal size). Dynamic processes were simulated based on 2730 myeloid progenitor cells^21^ during lineage commitment (S2), harboring three groups with similar cell proportions that represent different states along the differentiation trajectory. The complexity levels of simulated data were tuned by decreasing the probability of a gene to be differentially expressed and by varying the dispersion across all genes. Considering these parameters as an ordered pair, we created four different levels of data complexity (L1-L4). The rationale behind the parameter choice was the creation of a baseline data set (L1), readily clusterable for most tools (reference *matchSCore*). Subsequently, we modified conditions, where L2 presented a reduced number of differentially expressed genes and L3/L4 showed higher gene dispersion at different proportions of group markers. In the research context, tools with broad application spectrum might be preferable, as an *a priori* estimation of sample complexity is difficult (even with expected outcomes). As expected, most clustering algorithms performed better in the L1 data set as indicated by the Fowlkes and Mallows index (FMI, **Online Methods**; S1: average FMI= 0.73; S2: average FMI = 0.58, **Fig. 2**). With increased complexity we observed strong differences in the performance within and between methods, indicating distinct optimal application scenarios for the tested methods.

**Figure 2.**
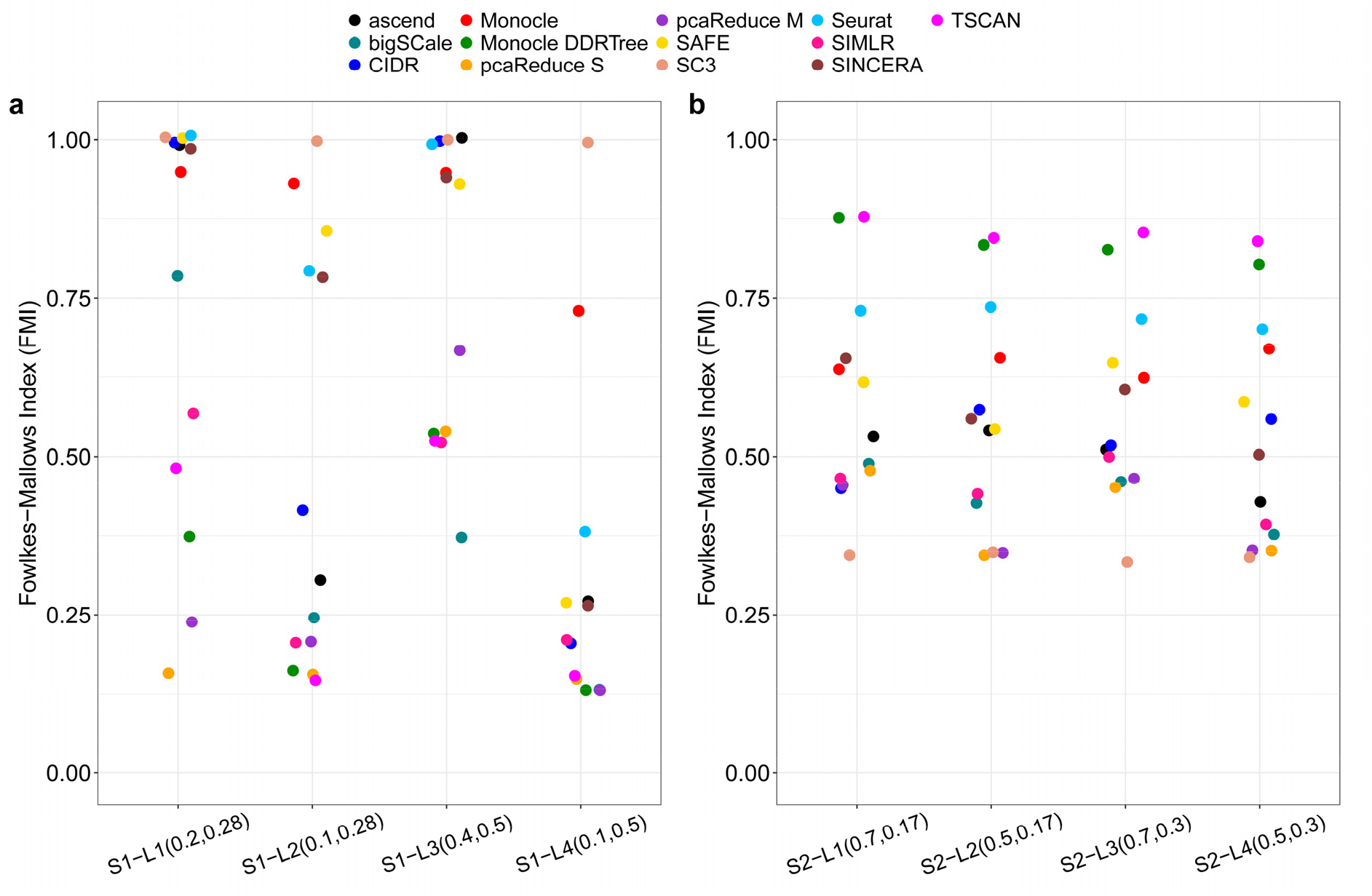
Distribution of the *Fowlkes-Mallows Index (FMI)* for each simulated data set, indicating the cluster accuracy for **a**) cell type (S1) and **b)** cell state (S2) scenarios. The x-axis labels indicate the level of complexity (L1-L4) with information about the simulation parameters (proportion of differentially expressed genes and gene dispersion, respectively). An FMI of 1 relates to complete agreement with true group labels, whereas 0 is comparable to a random assignation.

### Scenario 1 (S1 - Cell type heterogeneity)

Testing the performance of tools on data sets simulating distinct cell types and subclusters revealed strong differences in both clustering performance (FMI, **Fig. 2a**) and total *matchSCore* values (**Fig. 3, Supplementary Figs. 1-3**). As aforementioned, the *matchSCore* is not strictly reflecting the clustering performance, exemplified by the near perfect cluster identification of SINCERA^22^ (L1, FMI = 0.99; **Fig. 2a**), but a low total score due to a reduced capability to identify the group markers (**Fig. 3a,c**). *Vice versa*, bigSCale^23^ was outperformed in the clustering but reached the highest *matchSCore* by identifying the largest number of markers. Importantly, *matchSCore* curves not only provide a quantitative measure, but also give a qualitative indication of the accuracy across methods. For instance, SC3^24^ and Seurat BM^15^ showed different *matchSCore* trends (**Supplementary Fig. 4**), with Seurat BM showing a performance peak at 50% specificity (top 50% of real group markers) and the top 1000 ranked genes (computed by tool) per group, while SC3 detected an increasing number of marker with increasing specificity and tested genes (top genes). The continuously improving *matchSCore* in SC3 indicates the presence of true positive markers also at higher ranked positions, though the tool showed the overall lowest number of total markers in all simulations (**Supplementary Figs. 1-3**). SC3 displayed increased sensitivity at high specificity level (**Supplementary Fig. 1b**), in line with the original application spectrum. However, markers shared between two or more groups are detected as unique group-specific markers, resulting in overall reduced *matchSCore* values.

**Figure 3.**
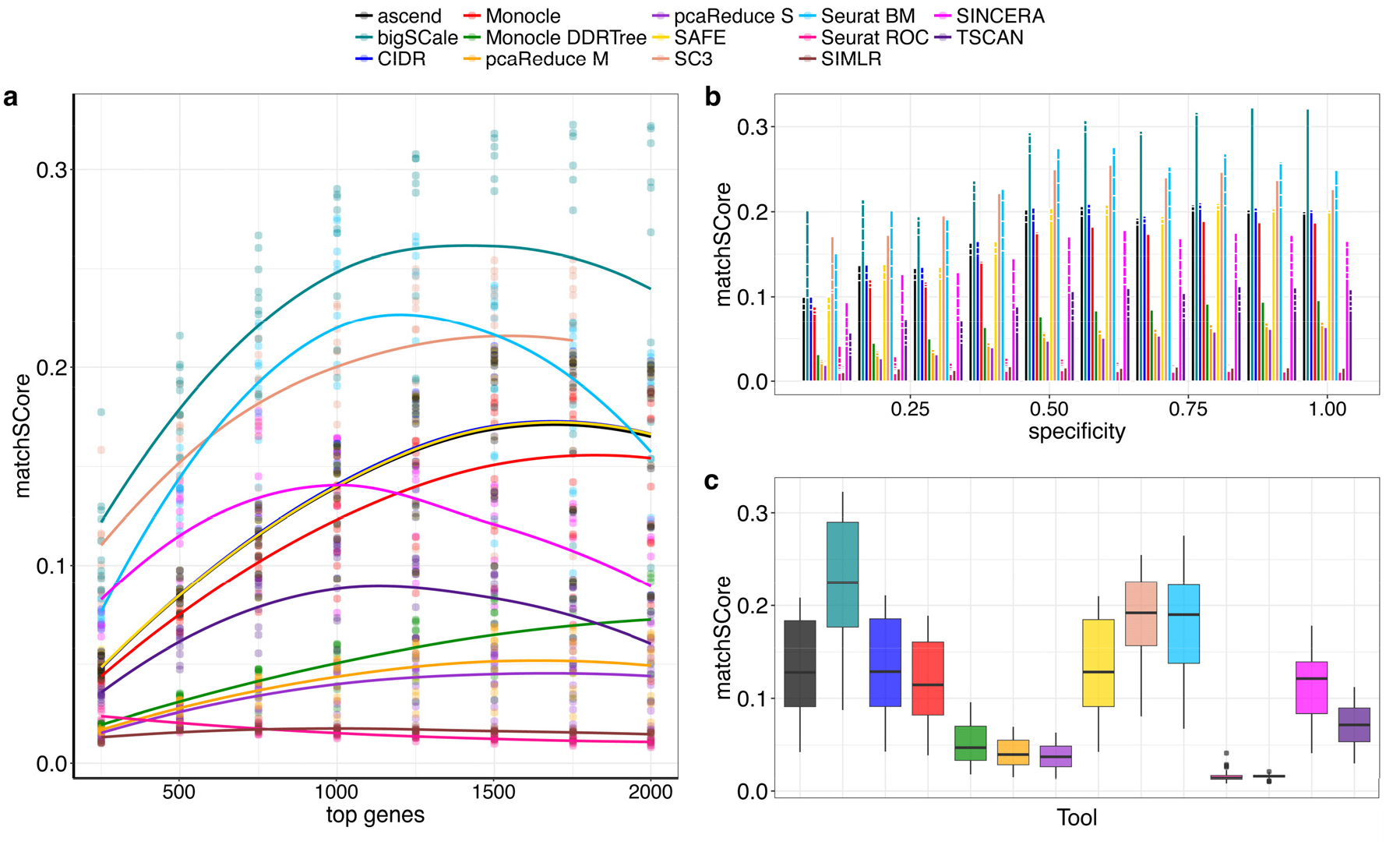
Benchmarking of 14 computational tools by using *matchSCore* for the simulation scenario 1 at level 1. *matchScores* are computed at different cutoffs (from 250 to 2000 by 250) of ranked markers (top genes). **a)** At each cutoff the plot shows the relative *matchSCore* at different level of specificity for the true group markers (represented by points). A local fitting regression (loess) was used to fit the trend of points. **b)** Barplots of *matchSCores* at different levels of specificity for group markers provided by the simulation. The value of specificity is indicated by the proportion of top ranked markers. **c)** Boxplots of *matchSCores* for all tested tools. Min, Max, 1^st^, 2^nd^ and 3^rd^ quartile are indicated considering all tested values of top-ranked markers.

At complexity level 2 (L2, **Supplementary Fig. 1**), the impact of the lower number of differentially expressed genes (7% of all genes compared to 14% in L1) seemed to split tool performance into two groups, with CIDR^25^, bigSCale, SIMLR^14^, Monocle DDRTree^16^, ascend^26^ and TSCAN^27^, producing low FMI values in clustering and also reduced *matchSCore* measures. In this context SC3 presented the highest *matchSCore (***Supplementary Fig. 1a,c**), considerably outperforming all other tools. The improved overall cluster performance in level 3 (L3, **Fig. 2a**) suggests that an increased number of differentially expressed genes (40% of total genes) allows the correct cluster assignation even with high dispersion. Here, SC3 and Seurat BM presented highest *matchSCore* values at lower numbers of top marker genes, which dropped with increasing marker quantity (**Supplementary Fig. 2)**. In contrast, bigSCale showed a continuously increasing trend, supporting its sensitivity in the detection of low-specificity markers. As expected, most of tools performed worse at complexity level 4 (L4, **Supplementary Fig. 3**), which included the highest gene dispersion and the lowest number of differentially expressed genes. Nevertheless, SC3 and Monocle clearly outperformed the other tools, with the former showing improved performance as the number of marker genes increase.

### Scenario 2 (S2 - Cell state dynamics)

In the second scenario simulating a continuum of cell states (instead of discrete cell types) we identified a strikingly different performance of many tools compared to the first scenario (**Supplementary Fig. 5-8**). In contrast with the analysis of simulated cell types, the analysis of cells in a dynamic context challenged the clustering and the assignment of cells to a unique group (**Fig. 2b**). Accordingly, none of the methods yielded perfect clustering. However, some approaches appeared more suitable in this context, especially tools developed to reconstruct continuous processes, such as Monocle DDRTree and TSCAN (average FMI > 0.8). It is important to note that low FMI values do not necessarily point to poor group assignment in the *matchSCore* cluster matching, if clusters are proportionally representing correct groups. However, the miss-assignment within clusters can impact on the marker identification, unless a tool is able to discriminate (giving different weights in the model) between good and poor representative cells. The fitted *matchSCore* curves resulted more flat in the second scenario, denoting a stability of *matchSCore* trends between high and low specificity markers in contrast with the previous scenarios.

At L1 the bigSCale *matchSCore* level largely outperformed all other tools (**Supplementary Fig. 5**), in contrast to its clustering capacity in this scenario (**Fig. 2b**). This is related to the increased presence of markers that are shared between the groups. Indeed, the dynamic nature of the simulated process was favorable for tools that better detect low-specificity markers, a strength of the bigSCale model. Next, TSCAN and Seurat BM had similar trends, although TSCAN resulted in overall higher *matchSCore* values. Both tools clearly differed from the other approaches in their accuracy in defining top-ranked gene markers. Noteworthy, Monocle DDRTree performance greatly improved when detecting less specific genes, while showing poor accuracy for the top-markers. At L2 (**Supplementary Fig. 6**), we observed a similar profile with stably good performance of TSCAN, Seurat BM and Monocle DDRTree. The bigSCale *matchSCore* decreased, due to the poorer clustering performance. L2 and L4 complexity simulations, as well as L1 and L3, had very similar *matchSCore* patterns supporting the correlation between the proportion of differentially expressed genes and tool performance. Overall, tools obtained higher *matchSCores* with more simulated differentially expressed genes, while at lower marker numbers, many methods reduced performance considerably. In all S2 simulations, Seurat ROC failed to identify markers, resulting in a constant *matchSCore* of zero. A more detailed interpretation of the tool benchmarking results is accessible in the **Supplementary Notes**.

### Mapping cell clusters across single-cell experiments

The logic behind deriving a *machSCore* does not only apply to the comparison of computational tools using a simulated ground truth, but also enables the matching of experimentally derived cell clusters to a reference data set. The fact that scores are computed over the reproducibility of population markers, equally allows matching of two different data sets and even across distinct scRNA-seq technologies. To formally test this assumption, we integrated experiments from human and mouse cell atlas studies that deconvoluted cell type composition of various organs by scRNA-seq. Specifically, we utilized Smart-seq2^28^ derived data sets as reference for cell type annotation and projected cell clusters from tissue-matched experiments onto this atlas. Smart-Seq2 was shown to produce high-quality single-cell expression profiles through the sensitive detection of RNA molecules, being most suitable to serve as reference technique^29, 30^.

As an illustrative example we selected the mouse bladder, an organ composed of two major cell types (luminal and mesenchymal cells) and a hierarchy of distinct subpopulations. Bladder tissues were included in two large-scale mouse atlas projects^2, 3^, providing a rich resource for data integration across experiments and technologies. Clustering of 1287 cells profiled using Smart-Seq2 led to the annotation of eight reference subpopulations (four mesenchymal, three luminal and one basal, **Fig. 4a,b**). When we then used *matchSCore* to integrate a microfluidic-based mouse bladder data set^2^ (Chromium, Single Cell 3’, **Fig. 4c,d**) we observed a clear consistency in cell type annotation between the test and reference data sets (**Fig. 4e**). Specifically, two mesenchymal subpopulations (A1 and A2) showed strong association to two reference populations of the same type, but being clearly distinct from other mesenchymal clusters (B1 and B2). In line, the luminal clusters A1 and B were related to two distinct reference subpopulations, while an intermediate cluster (A2) presented association to all luminal types. The expression of population specific signatures (**Fig. 4f**) underscored the correct cluster classification using *matchSCore*. Noteworthy, immune and endothelial cells that have exclusively been included in the test experiment were not assigned to any reference subpopulations (**Fig. 4e**). When we integrated a second data set derived from mouse bladder that has not been annotated before (Microwell-Seq^3^; cluster 1-16), we could associate all reference subpopulations to specific clusters (**Supplementary Fig. 9**). Gene markers for unassigned clusters did not show an enrichment in any of the reference subpopulations, suggesting the absence of the cell types in the reference data set (e.g. blood cell types).

**Figure 4.**
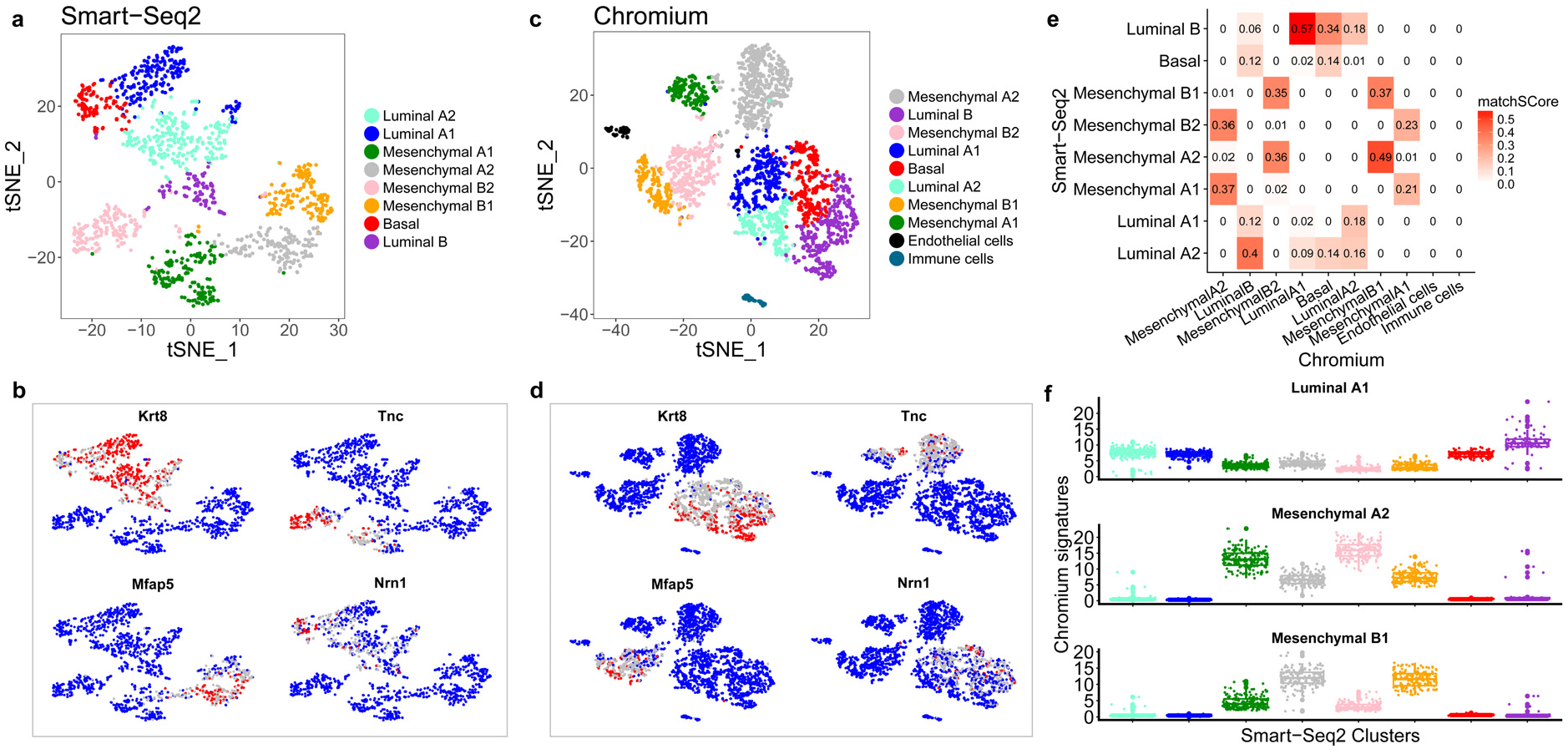
Data integration of bladder samples from two different experiments and technologies (Smart-seq2 and Chromium). **a,c)** The t-SNE plot displays the different cell populations identified by the clustering of the **a)** Smart-Seq2 (reference data) and **c)** Chromium (test data) experiment^2^. **b,d)** t-SNE plots showing the relative expression of most significant bladder markers (blue: low; red: high; grey: intermediate) for **b)** Smart-Seq2 and **d)** Chromium data sets. **e)** *matchSCore* values computed by comparing the test clusters (Chromium) against the reference cell groups (Smart-Seq2). **f)** Relative expression of the matching cluster signatures (top 100 ranked genes) from the test data within cell reference populations.

To further confirm the applicability of *matchSCore* guided data integration, we produced a single-cell RNA sequencing data set for mouse bladder cells using MARS-seq^31^. Specifically, we isolated urothelial epithelial cells from normal mouse bladder by FACS by negative selection of leukocytes (CD45), fibroblasts (CD140a), endothelial cells (CD31), and erythrocytes (Ter119) and derived scRNA-seq data for 1018 cells. Cell clustering identified four putative subpopulations and respective gene markers. To annotate these experimentally derived populations, we used *matchSCore* to integrate our results with the Smart-Seq2 reference data set (**Supplementary 10a,b**). Two principal clusters could be clearly assigned to luminal and mesenchymal reference cell types of the bladder; and two additional subclusters showed enrichment in luminal subpopulations. Consistently, cells assigned to luminal populations (cluster 0,1,2) presented high levels of related signatures and the cell type specific markers *Krt5 and Upk1a*; while mesenchymal signatures and markers (*Vim* and *Upk3b*) were specific for cluster 3 (**Supplementary Fig. 10c,d**).

Considering the wealth of reference data sets that are being created within the framework of the Human Cell Atlas project^1^, we further aimed to confirm the suitability of *matchSCore* annotation in a human context. Such integration tool will be extremely powerful to interpret future experiments in respect to a predefined human reference atlas. We used human pancreas, an organ with a well-known anatomy and defined cellular subtypes. As before, we used an annotated Smart-Seq2 data set^2^ as reference before matching cell cluster from different studies and techniques. The reference atlas (**Supplementary Fig. 11**) described five subpopulations of endocrine pancreatic islets cells (PP, alpha, beta 1/2 and delta cells) and five additional cell types (stellate, ductal, exocrine, endothelial and immune). We projected two well-annotated (CEL-seq^32^, **Supplementary Fig. 12a**; Indrop^33^, **Supplementary Fig. 13a**) and an unannotated (Microwell-Seq^3^; **Supplementary Fig. 14a**) pancreas data sets on the reference map and could assign clusters to all reference subpopulations. Population specific signatures supported the correct annotation of the clusters and the accuracy of our approach (**Supplementary Fig. 12b,13b,14b**). Of note, stellate cells were assigned to multiple clusters, likely representing different activation states (activated or quiescent^33^, **Supplementary Figs. 13a** and **14a**). Also immune cells had multiple matches, pointing to the different blood cell types. To further validate the broad utility and sensitivity of our approach in a human context we produced and projected MARS-Seq data from 289 short-term cultured gradient-purified primary human pancreatic islet cells (**Fig. 5a,b**). Consistently, the largest cell clusters (Cluster_0 and Cluster_1) matched to endocrine subpopulations, with scores and population signatures distinguishing alpha and beta cell types (**Fig. 5c,d**). In addition, minor populations of contaminating duct and stellate cells could be clearly annotated.

**Figure 5.**
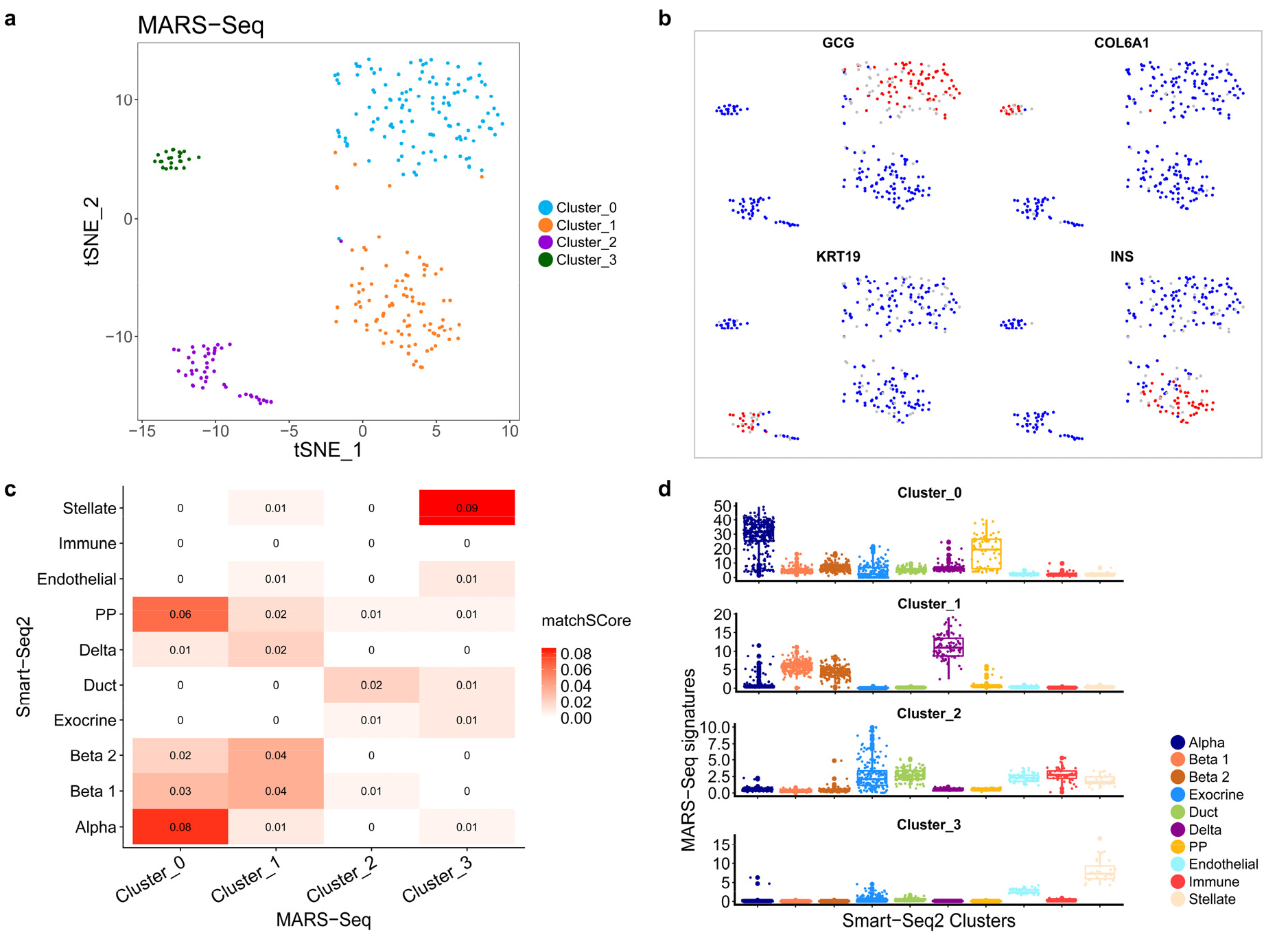
Using *matchSCore* to annotate pancreatic islet cell types derived from MARS-Seq. **a)** The t-SNE plot displays the different cell populations identified by the clustering analysis of 289 pancreas cells from MARS-Seq. **b)** t-SNE plots displaying the expression (blue: low; red: high; grey: intermediate) of pancreatic subpopulation markers. **c)** *matchSCore* values computed by comparing the test clusters (MARS-Seq) against the reference cell groups (Smart-Seq2). Each tested cluster is matched against all reference clusters in order to find its most similar reference group. **d)** Relative expression of the matching cluster signatures (top 100 ranked genes) from the MARS-seq experiment within the reference clusters (Smart-Seq2).

Conclusively, the organ-matched integration of six different RNA sequencing technologies and nine experiments using *matchSCore* pointed to the conservation of sufficient gene marker signal across methods and varying sequencing depths. Moreover, it illustrates the broad application spectrum of *matchSCore* to integrate cell populations from different experiments.

## Discussion

Cellular phenotyping using RNA sequencing is at the forefront at describing cell types and states. The resolution to which a sample is characterized relates to technical features, such as library preparation method and sequencing depth, but also to the choice of data analytic strategy. Computational tools are conceptually different and harbor largely different statistical approaches that impact on their sensitivity to describe cellular phenotypes. In general, cell clustering approaches detect patterns in a data set and group cells that co-express gene sets. However, different clustering algorithms prioritize genes differently and, thus, can come to different conclusions. We developed a scoring system that allowed us to systematically test single-cell phenotyping tools for their accuracy and sensitivity to chart sample heterogeneity by comparing them against a simulated ground truth. We were able to quantify the combined performance of clustering and marker detection at different levels of specificity and two biological single-cell scenarios. Consistent with the fact that tool design centered on cell clustering, most tools showed decent accuracy in group assignments, but often failed to determine cluster-specific markers. In addition, tools have different capacity to deal with more challenging data types, exemplified by dynamic systems. **Figure 6** summarizes the performance for both biological scenarios and the different levels of complexity. Here, we display the relative average *matchSCore* (across different levels of top-markers) as a measure of accuracy in retrieving a cell’s phenotype.

**Figure 6.**
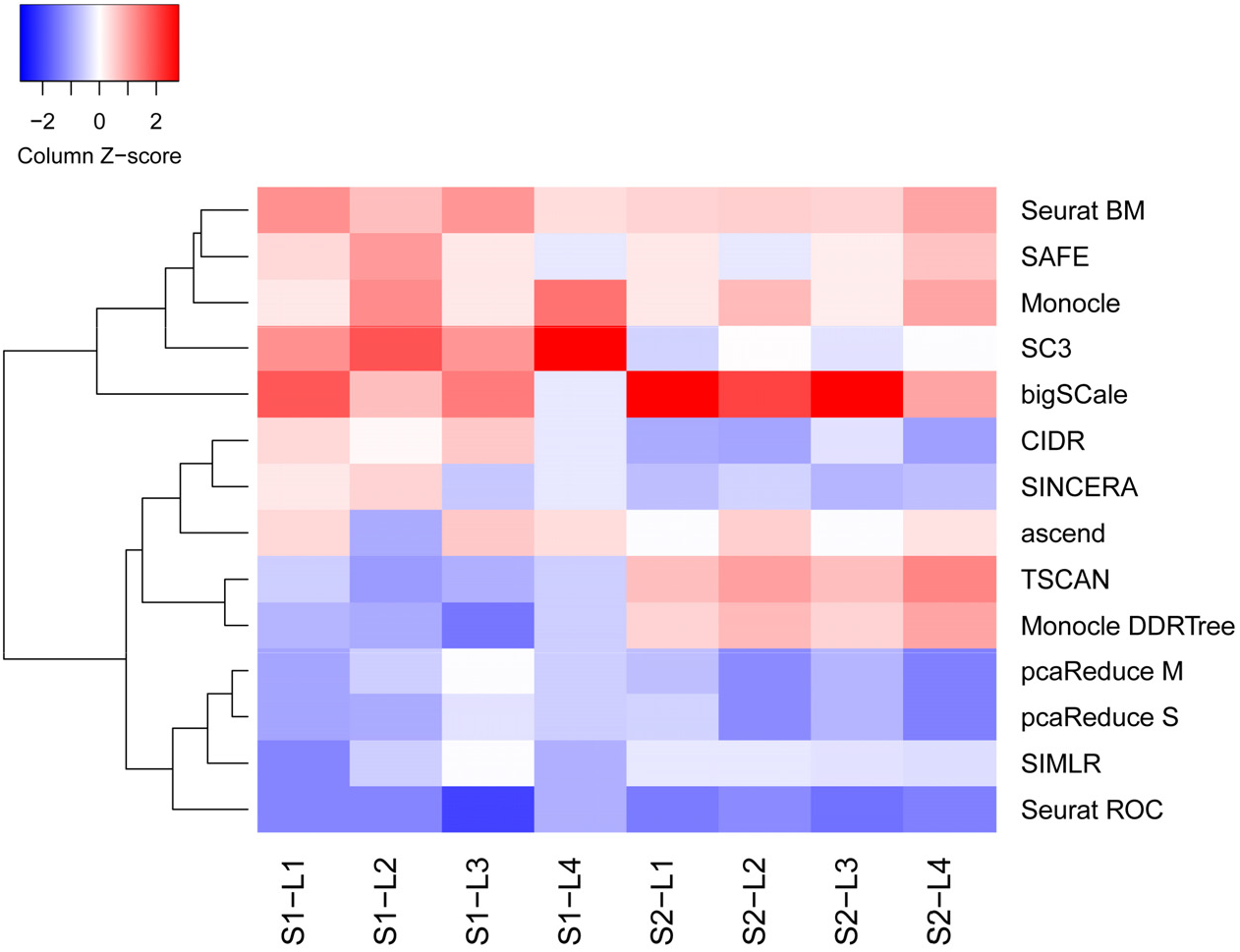
Summary heatmap of the performance of computational tools. For each scenario and level of complexity, the normalized average across different thresholds of top markers (from 250 to 2000 by 250) *matchSCores* are shown as Z-scores. Higher Z-scores are related to better performance.

The logic in our *matchSCore* system also allowed us to enlarge its application to the comparative analysis of phenotypes across data sets, thus providing a straightforward solution to annotate future single-cell projects. Clustering single cells into a subtype structure provides unprecedented resolution of complex samples, however, data interpretation is extremely challenging. Here, the data-driven results often exceed prior knowledge about tissue composition and novel cell clusters are not straightforward to characterize. Consequently, comparative analysis between experiments can guide researchers to interpret their data in respect to related experiments or previously produced references. *matchSCore* enables this straightforward cross-annotation between studies, even in cases of data types from largely different scRNA-seq methodologies. We have provided examples for human and mouse tissue atlas projects and selected organs. However, our method is readily applicable for any single-cell research context and species. Compared to other strategies to project clusters into reference maps, *matchSCore* can be directly applied at the gene marker level, thereby combining findings (post clustering) rather than through data integration. In fact, published integration approaches using nomalization are combining data in its rawest format without providing a straighforward solution to compare cell identities^17, 19^. Notably, our approach showed to be also viable when reference and test samples included non-overlapping populations, which can lead to misinterpretations particularly when rare subpopulations are present. A recent approach for direct cell projection^18^ includes several heuristic steps and arbitrary cutoffs, which could amplify technical biases, reducing the power of classification. The approach also makes use of distance metrics (Pearson, Spearman and Cosine) that are not tailored to the exceptionally sparse and noisy nature of scRNA-seq data sets, which could lead to shortcomings during data interpretation.

In conclusion, *matchSCore* provides a direct and reliable solution to match cell identities across single-cell analyses and data sets, allowing to perform a systematic comparison of computational and experimental outcomes, respectively. Considering the rapid computational tool development and large-scale data production, *matchSCore* can contribute to define high-quality analysis standards and the meaningful data interpretation.

## Acknowledgements

We thank A. Lafzi and G. Esteban-Rodriguez for support and discussion; and L. Martínez for help with flow cytometry analyses. HH is a Miguel Servet (CP14/00229) researcher funded by the Spanish Institute of Health Carlos III (ISCIII). This work was funded, in part, by a grant from the Fundación Científica de la Asociación Española Contra el Cáncer /AECC) to FXR. Core funding is from the ISCIII and the Generalitat de Catalunya.

## Author contributions

HH and EM conceived the study. EM developed matchSCore and performed the statistical analysis. GI performed bigSCale analyses. AGA, CM and GL generated sequencing libraries and helped with the data interpretation. JF, IMR, FR and CPS contributed human pancreas and mouse bladder samples. IG provided the computing infrastructure. HH and EM wrote the manuscript. All authors read and approved the final manuscript.

## Competing financial interests

The authors declare no competing financial interests.

## Availability of the source code

All functions of *matchSCore* are available at Github under the link: https://github.com/elimereu/matchSCore

## Online Methods

### The matchSCore metric

The *matchSCore* quantifies the consistency in the assignment of cell clusters and gene marker across analyses and experiments. Specifically, it measures the combined accuracy of clustering and identification of cluster-specific markers with respect to the optimal solution, provided by the simulation. In addition, it enables to match ranked marker genes between experiments and to convert their agreement in a score. To benchmark analysis tools, a matching step between clusters and simulated groups was performed to assign true group labels to each cluster. The label for each cluster is defined according to the most frequent original cell group present in a given cluster. Once labels have been assigned to clusters, a Jaccard Index is used to measure similarities across corresponding cluster markers and true (simulated) group markers. The Jaccard index is commonly used to quantify the similarity of two sample sets by computing their intersection over their union. For shared markers across groups, the Jaccard index allows to have elements in more than one group, an important feature in the context of cellular phenotyping. In order to compare different experiments, cluster markers were matched against all group markers from the reference sample and a *matchSCore* was computed for all combinations. Group markers were ranked according to their specificity and the 100 top-ranked markers were used for all data set comparisons.

### Simulated data

To simulate data we used Splatter^34^, which allows to estimate parameters from real data sets and to control the proportion of differentially expressed genes and dispersion. When simulating several groups or paths, Splatter stores information about group fold-changes for each gene, which is needed for the ranking of group-specific markers. The complexity levels of our simulated data sets were defined by two parameters: the probability of a gene to be differentially expressed (de.prob) and the underlying common dispersion across genes (cv). Considering these parameters as an ordered pair (de.prob,cv), we created four different data quality levels L1, L2, L3 and L4 with the corresponding pairs (0.2, 0.28), (0.1,0.28), (0.4,0.5), (0.1,0.5) in scenario 1 and (0.7,0.17), (0.5,0.17), (0.7,0.3), (0.5,0.3) in scenario 2. The rationale behind the choice of combinations was to determine a reference *matchSCore* with a baseline data set (L1), for which most of tools perform well at clustering level. In subsequent levels we then increased the complexity with L2 presenting a reduced number of differentially expressed genes and L3 and L4 higher gene dispersion at different proportions of differentially expressed genes.

### Definition of gene markers and their ranking

The definition of a cell type is strictly related to the markers used for their characterization. A cell type marker is a gene that is positively and stably differentially expressed in a given group with respect to a reference gene mean. However, in order to have a more complete insight into complex samples and dynamic process, shared markers between cell types or states can also be relevant. Accordingly, we ranked markers based on their specificity in groups, where lower ranks reflect higher specificity level and higher ranking positions include genes shared across groups. Here, the sorting of markers was performed according to their fold-changes, with higher fold-changes ranking at top positions. For each of the tested tools, cluster-specific ranking has been performed ordering p-values by significance, resulting in a ranked list of genes for each cluster.

With the aim to quantify the accuracy of methods in ranking group markers, we set different thresholds of specificity. Specifically, we define a common proportion of top ranked markers across all groups before comparing the simulated true markers with those predicted by the methods (at different numbers (k) of top genes). For example, from the simulation point of view a specificity of 0.1 relates to the top 10% markers per group. Consequently, tools reaching higher *matchSCore* values at high specificity levels (low percentages of top markers) are more accurate in the detection of markers specific for each group. For lower specificities, *matchSCores* will progressively increase with the predicted matching the true markers until a maximum is reached. Best performing tools show an increasing *matchSCore* function as the number of predicted markers get larger, indicating that they are able to detect not only cluster-specific but also higher order cluster markers.

### Clustering and identification of cluster-specific markers

The benchmarking involved 14 methods from 11 different tools. Some tools provide more than one approach for clustering (Monocle, pcaReduce, Seurat) and for the identification of markers (Seurat). Despite the large number of tools for clustering scRNA-seq data, only few include the detection of cluster-specific markers. Consequently, for those tools that do not provide the computation of cluster-specific markers, we used MAST^35^ (as implemented in the Seurat package), a flexible and scalable tool for differentially expression analysis and specifically designed for single-cell data. Before conducting MAST, log-normalization and scaling has been applied for tools that do not provide a normalized count matrix. The cluster accuracy has been evaluated by the Fowlkes and Mallows Index (FMI), a metric defined as the geometric mean of precision and recall. The index ranges between 0 and 1, where 1 represents perfect match of the two partitions and 0 completely independent clustering. The parameters of all methods were set to their default setting, but we used the exact number of clusters as defined in the simulated data set. All data sets and cell type annotations were downloaded from their public accessions. Clustering (if missing) and marker identification were done using Seurat with default settings.

### Cell isolation from primary mouse bladder and human pancreas tissues

Mice were sacrificed and the bladder was accessed and turned inside-out leaving the urothelial surface exposed. The urothelium was enzymatically digested with collagenase P (0.5μg/mL) in Hank’s Balanced Salt Solution (HBSS) in a thermoblock with gentle shaking at 37°C for 20min. Collagenase P was inactivated with 2mM EDTA and 50% of fetal bovine serum. The cell suspension was collected and the remaining urothelium was scraped. After filtering through a 70μm strainer and centrifugation at 1200rpm for 5min at 4°C, cells were washed 1x in PBS and incubated with blocking buffer (1% BSA/3mM EDTA in PBS) for 15min at room temperature. After washing 2x with PBS, cells were incubated with APC-labeled anti-CD45 (BD biosciences, Cat. No. 559864), PE-labeled anti-CD31 (BD biosciences, Cat. No. 555027), PE-labeled anti-CD140a (Labclinics, Cat. No. 16-1401-82) and PE-labeled anti-Ter119 (BD biosciences, Cat. No. 116208) antibodies in FACS buffer (0.1% BSA / 3mM EDTA in PBS) for 30min at 4°C. After washing 2x with PBS, cells were resuspended in FACS buffer and stained with DAPI (Sigma-Aldrich). A control sample lacking primary antibody and a Fluorescence Minus One (FMO) control were used in all experiments. All samples were analysed using a FACS Influx or AriaII (BD Biosciences) flow cytometer and at least 10,000 events were acquired. Analyses were performed using FlowJo flow cytometer analysis software.

Human pancreatic islets from organ donors were isolated and purified using established isolation procedures previously described^36^, shipped in culture medium and then re-cultured at 37°C in a humidified chamber with 5% CO2 in RPMI 1640 medium supplemented with 10% fetal calf serum, 100 U/ml penicillin, and 100 U/ml streptomycin for three days prior to dissociation and FACS isolation.

### Library preparation and sequencing

To construct single-cell libraries from poly(A)-tailed RNA, we applied massively parallel single-cell RNA sequencing (MARS-Seq)^21, 37^. Briefly, single cells were FACS isolated into 384-well plates, containing lysis buffer (0.2% Triton X-100 (Sigma-Aldrich); RNase inhibitor (Invitrogen)) and reverse-transcription (RT) primers. The RT primers contained the single-cell barcodes and unique molecular identifiers (UMIs) for subsequent de-multiplexing and correction for amplification biases, respectively. Single-cell lysates were denatured and immediately placed on ice. The RT reaction mix, containing SuperScript III reverse transcriptase (Invitrogen) was added to each sample. After RT, the cDNA was pooled using an automated pipeline (epMotion, Eppendorf). Unbound primers were eliminated by incubating the cDNA with exonuclease I (NEB). A second pooling was performed through cleanup with SPRI magnetic beads (Beckman Coulter). Subsequently, pooled cDNAs were converted into double-stranded DNA with the Second Strand Synthesis enzyme (NEB), followed by clean-up and linear amplification by T7 *in vitro* transcription overnight. Afterwards, the DNA template was removed by Turbo DNase I (Ambion) and the RNA was purified with SPRI beads. Amplified RNA was chemically fragmented with Zn2+ (Ambion), then purified with SPRI beads. The fragmented RNA was ligated with ligation primers containing a pool barcode and partial Illumina Read1 sequencing adapter using T4 RNA ligase I (NEB). Ligated products were reverse-transcribed using the Affinity Script RT enzyme (Agilent Technologies) and a primer complementary to the ligated adapter, partial Read1. The cDNA was purified with SPRI beads. Libraries were completed through a PCR step using the KAPA Hifi Hotstart ReadyMix (Kapa Biosystems) and a forward primer that contains Illumina P5-Read1 sequence and the reverse primer containing the P7-Read2 sequence. The final library was purified with SPRI beads to remove excess primers. Library concentration and molecular size were determined with High Sensitivity DNA Chip (Agilent Technologies). The libraries consisted of 192 single-cell pools. Multiplexed pools were run on Illumina HiSeq 2500 Rapid flow cells following the manufacturer’s protocol. Primary data analysis was carried out with the standard Illumina pipeline. We produced 52 nt of transcript sequence reads.

### MARS-Seq data processing

The MARS-Seq technique takes advantage of two-level indexing that allows the multiplexed sequencing of 192 cells per pool and multiple pools per sequencing lane. Sequencing was carried out as paired-end reads; wherein the first read contains the transcript sequence and the second read the cell barcode and UMI. Quality check of the generated reads was performed with the FastQC quality control suite. Samples that reached the quality standards were then processed to deconvolute the reads to single-cell level by de-multiplexing according to the cell and pool barcodes. Reads were filtered to remove poly(T) sequences. Reads were mapped with the RNA pipeline of the GEMTools 1.7.0 suite^38^ using default parameters (6% of mismatches, minimum of 80% matched bases, and minimum quality threshold of 26) and the genome references for mouse (Gencode release M15, assembly GRCm38) or human (Gencode release 26, assembly GRCh38). Gene quantification was performed using UMI corrected transcript information to correct for amplification biases, collapsing read counts for reads mapping on a gene with the same UMI (allowing an edit distance up to 2 nt in UMI comparisons). Only unambiguously mapped reads were considered. Before clustering low quality cells were filtered out by removing cells having higher mapping rate to ERCC spike-ins and mitochondrial genes. Further, we removed cells with a log number of detected genes lower than expected (more than 3 median absolute deviations below the median). In addition, we positively filtered for genes detected in at least 5 cells, having a minimum level of expression greater or equal to the median expression value across all genes.

